# Inhibition of CRISPR-Cas12a DNA Targeting by Nucleosomes and Chromatin

**DOI:** 10.1101/2020.07.18.210054

**Authors:** Isabel Strohkendl, Fatema A. Saifuddin, Bryan A. Gibson, Michael K. Rosen, Rick Russell, Ilya J. Finkelstein

**Affiliations:** Department of Molecular Biosciences and Institute for Cellular and Molecular Biology, University of Texas at Austin, Austin, Texas 78712, USA; Center for Systems and Synthetic Biology, University of Texas at Austin, Austin, Texas 78712, USA; Department of Biophysics and Howard Hughes Medical Institute, University of Texas Southwestern Medical Center, Dallas, TX 75390, USA

**Keywords:** Cpf1, kinetics, rate-limiting binding, nucleosome arrays, liquid-liquid phase separation

## Abstract

Genome engineering nucleases, including CRISPR-Cas12a, must access chromatinized DNA. Here, we investigate how *Acidaminococcus sp*. Cas12a cleaves DNA within human nucleosomes and phase-condensed nucleosome arrays. Using quantitative kinetics approaches, we show that dynamic nucleosome unwrapping regulates DNA target accessibility to Cas12a. Nucleosome unwrapping determines the extent to which both steps of Cas12a binding–PAM recognition and R-loop formation–are inhibited by the nucleosome. Nucleosomes inhibit Cas12a binding even beyond the canonical core particle. Relaxing DNA wrapping within the nucleosome by reducing DNA bendability, adding histone modifications, or introducing a target-proximal nuclease-inactive Cas9 enhances DNA cleavage rates over 10-fold. Surprisingly, Cas12a readily cleaves DNA linking nucleosomes within chromatin-like phase separated nucleosome arrays—with DNA targeting reduced only ~4-fold. This work provides a mechanism for the observation that on-target cleavage within nucleosomes occurs less often than off-target cleavage within nucleosome-depleted regions of cells. We conclude that nucleosome wrapping restricts accessibility to CRISPR-Cas nucleases and anticipate that increasing nucleosome breathing dynamics will improve DNA binding and cleavage in eukaryotic cells.

## Introduction

Class 2 CRISPR-Cas nucleases Cas12a and Cas9 are precision genome editing tools that are evolutionarily and structurally distinct yet share several key reaction steps^1–3^. Class 2 nucleases first identify a protospacer adjacent motif (PAM) via protein-DNA contacts. PAM binding causes local DNA unwinding and then initiates directional strand invasion by the CRISPR RNA (crRNA)^4^. An R-loop between the crRNA guide sequence and the complementary target DNA extends towards the PAM-distal end of the target DNA strand. Once the R-loop is fully formed, the nuclease is activated and cleaves the target and non-target DNA strands to generate a DNA break^5^. R-loop extension is rate-limiting for cleavage by Cas12a and Cas9 *in vitro*^6–9^, providing a mechanistic explanation for Cas12a’s observed higher specificity *in vivo*^8^ ^10–15^.

In eukaryotic cells Cas nucleases must navigate chromatin, yet we only have a limited understanding of how chromatin reduces nuclease activity. To date, all investigations of Cas nuclease activity on chromatin have focused on Cas9-induced gene editing, and none on Cas12a. Cas9 activity is strongly dependent on the chromatin state, with actively transcribed regions being more susceptible to cleavage than dense heterochromatin^16–21^. Cas9 cleavage is depleted at nucleosomal sites in cells^21, 22^ and Cas9 cleavage *in vitro* shows near-complete inhibition approaching the nucleosome dyad^21, 23, 24^. While these studies have established that Cas9 is inhibited by nucleosomes, we do not fully understand the kinetic basis for this inhibition nor how these mechanistic results extend to Cas12a.

Nucleosomes are the smallest structural unit of the eukaryotic genome organization, wrapping 147 base pairs (bp) of DNA ~1.7 times around a histone octamer^25–27^. Consecutive nucleosomes are separated by 10-50 bp of linker DNA that appear as beads on a string^28–31^. Textbook models of higher order chromatin organization picture nucleosomes arranged into a 30-nm fiber, whereas recent findings suggest that chromatin is less structured^32^. Individual nucleosomes are dynamic assemblies^32^ in which DNA transiently unwraps from the histone octamer and rewraps in a phenomenon known as ‘nucleosome breathing’^33, 34^. Transcription factors and other site-specific DNA-binding proteins exploit nucleosome breathing to bind their target sites^35, 36^, but the stepwise unwrapping of DNA from nucleosome edges leads to reduced site exposure approaching the nucleosome dyad^37, 38^. Nucleosomes also increase the dissociation of transcription factors by several orders of magnitude^39^. The dynamics and accessibility of nucleosomal DNA may be further impacted by electrostatic interactions between the highly polarized nucleosomes^40^. Indeed, the high valence of nucleosome arrays enables ion-driven self-association into condensed, phase-separated droplets, mimicking chromatin in the nucleus^41, 42^. Mechanistically defining the impact chromatin has on Cas12a DNA targeting could help develop strategies for improving the efficiency and specificity of Cas12a as a genome editing tool.

Here, we analyze Cas12a cleavage kinetics on nucleosomal substrates. The unwrapping dynamics of a mono nucleosome regulate Cas12a cleavage efficiency, as was previously reported for Cas9^23^. Our data are consistent with a model in which nucleosome breathing regulates accessibility to Cas12a binding^36, 43, 44^. At nucleosome edges, both PAM recognition and R-loop propagation by Cas12a are inhibited, with an overall binding inhibition of several orders of magnitude. Surprisingly, we see significant cleavage inhibition up to 12 bp beyond the canonical core particle. Unwrapping the nucleosome by DNA sequence changes, histone modifications, or nuclease-dead Cas9 (dCas9) binding increases access of Cas12a. In contrast, the linker sequences between histones remain readily accessible in higher-order phase-separated assemblies representative of chromatin. Our results define the impact of nucleosomes on Cas12a activity and highlight strategies for modulating nucleosome breathing dynamics to increase DNA accessibility in eukaryotic genome editing.

## Results

### Cas12a cleavage is Inhibited Across the Entire Nucleosome Wrap

To understand how the nucleosome influences Cas12a cleavage, we designed crRNAs to target sequences adjacent to 5’-TTTA PAMs within an extended ‘Widom 601’ nucleosome positioning DNA sequence (**Figure 1A**, **Tables S1**, **S2**)^45^. The DNA was extended to include two Cas12a targets that are 16 and 34 bp beyond the nucleosome core particle. Using a strong positioning sequence permits efficient reconstitution of homogenous nucleosomes and nucleosome arrays (see below) and allows direct comparisons with other studies conducted on this DNA substrate^21, 23, 24, 46, 47^. A fluorescent end label was introduced into the DNA substrate during PCR amplification. Human nucleosomes were reconstituted via salt dialysis of the DNA with the purified human histone octamer^48^ (**Figure S1A**). Purified *Acidaminococcus sp* Cas12a (henceforth referred to as Cas12a; **Figure S1B**) was assembled with precursor crRNA into a ribonucleoprotein (RNP) complex that was directed to a single target in the DNA substrate. Cleavage reactions for each Cas12a target were carried out in parallel on both the DNA and nucleosome substrates at 25 °C. Time points were sampled from the reaction and stopped by treatment with EDTA and Proteinase K to remove bound histones and Cas12a, resolved on native polyacrylamide gels, and quantified to obtain the cleavage rates (**Figure 1B**, **Figure S1C**, **Table S4**).

**Figure 1.**
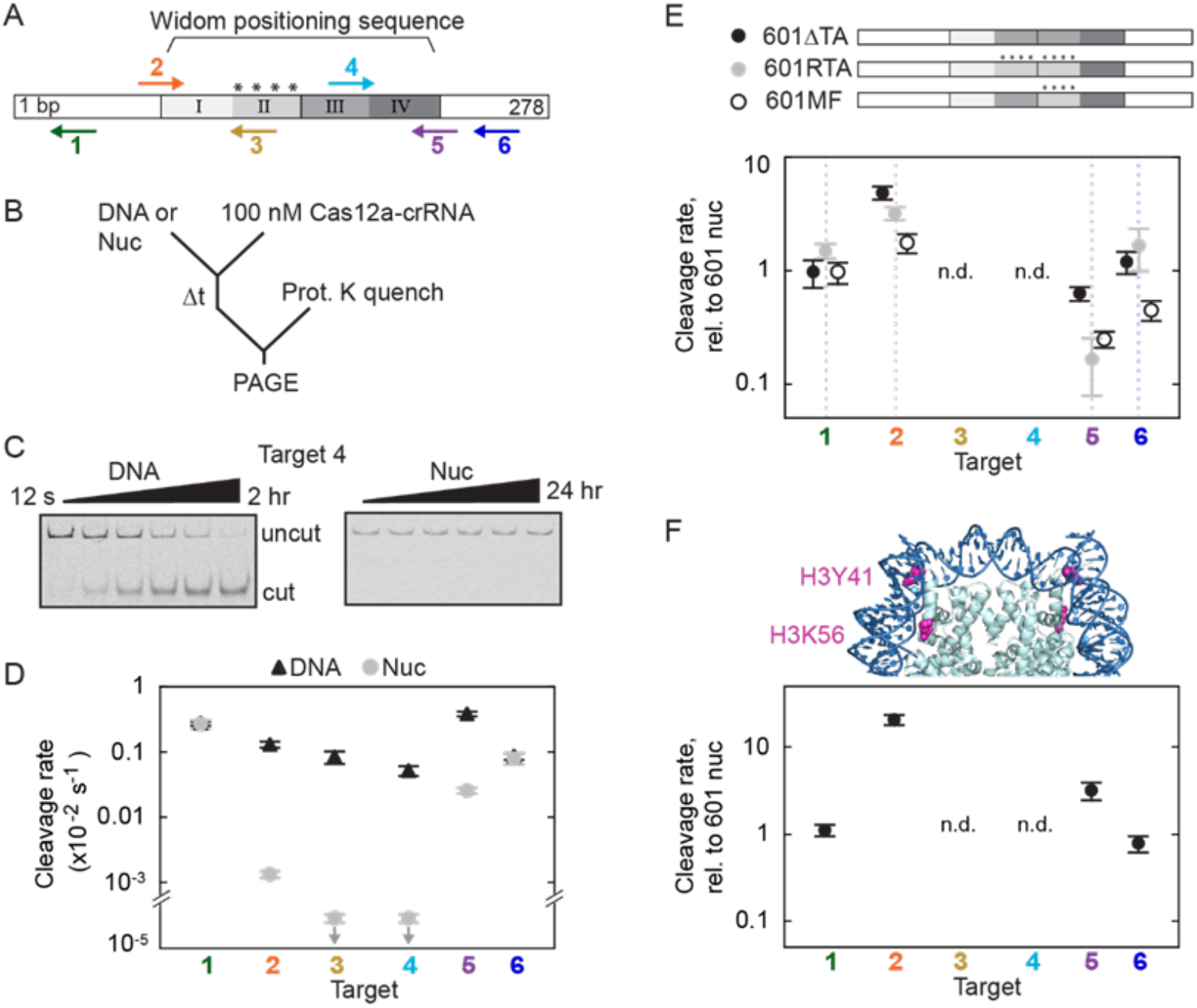
DNA unwrapping regulates Casl2a cleavage of nucleosomal targets. A) Diagram of the DNA substrate used in Cas12a cleavage reactions. The Widom 601 positioning sequence is divided into quartiles that indicate the inner (II, III) and outer (I, IV) wrap. Asterisks: the four TA dinucleotide repeat sets that produce tight wrapping of the inner left quartile. Arrows: Cas12a targets point in the direction of R-loop formation. For the top arrows, an R-loop forms with the Crick (bottom) strand and for bottom arrows, the R-loop forms with the Watson (top) strand. B) Cleavage reaction setup. ‘Nuc’: nucleosome substrate. C) Gels showing Cas12a-mediated cleavage of target 4. D) Plotted cleavage rates of the six Cas12a targets for DNA and nucleosome substrates at 25 °C. Downward arrows signify that the value is an upper limit estimated from the lack of detectable cleavage. E) Top: diagram depicting variant 601 constructs. Bottom: Cas12a cleavage of variant nucleosome substrates normalized to original 601nucleosome substrate rates. F) Top: crystal structure of a 601 nucleosome (PDB: 3LZ0^81^) with the two modified amino acids highlighted in magenta. H3Y41E and H3K56Q mimic H3Y41 ph and H3K56ac. Bottom: Relative cleavage of nucleosomal DNA reconstituted with H3(Y41 E/K56Q). D-F) Each data point is the mean of at least three replicates. Error bars: standard error of the mean (SEM). E and F) ‘n.d.’: ‘no data’ as no nucleosomal cleavage was observed for targets 3 and 4 for all nucleosome substrates.

Cas12a cleavage was inhibited for all nucleosome-occluded targets. Targets 3 and 4, positioned within the central wrap of the nucleosome, showed no detectable cleavage after 24 hrs (<6 ×10^−7^ s^−1^, **Figure 1C**), indicating inhibition of at least 1,000-fold relative to the nucleosome-free DNA. (**Figure 1C, 1D**). Targets 2 and 5, only partially overlapping with the edges of the nucleosome, were inhibited 100- and 15-fold respectively (target 2: 1.35 (±0.16) ×10^−5^ s^−1^; target 5: 2.58 (±0.29) ×10^−4^ s^−1^, mean ± SEM of three biological replicates). We attribute the asymmetric inhibition of the edge targets to the non-palindromic 601 sequence and its asymmetric interactions with the histone octamer. The higher inhibition observed for target 2 is consistent with prior biophysical studies that noted greater bendability and tighter wrapping of the DNA around the histone octamer at this site (also see below)^46, 49^. Targets 1 and 6, positioned in the nucleosome-flanking DNA, were cleaved comparably to nucleosome-free DNA substrates with rates of 2.72 (±0.40) ×10^−3^ s^−1^ and 8.2 (±1.6) ×10^−4^ s^−1^, respectively. Interestingly, the cleavage rates of the targets on the DNA substrates varied six-fold. We attribute this variation to DNA sequence-dependent binding and cleavage (**Figure 1D**). In sum, our results show strong position-dependent cleavage inhibition by the nucleosome.

We next measured nucleosome-mediated inhibition of Cas12a at 37 °C (**Figure S1D**). Cleavage rates of the isolated DNA increased up to 13-fold and maintained a similar pattern of preferred target sequences, although now within a smaller range of ~3-fold. In the context of the nucleosome, the inner wrap targets 3 and 4 showed detectable cleavage after 24 hours, resulting in ~2,000-fold inhibition with rates of 5.2 (±1.9) ×10^−6^ s^−1^ and 4.2 (±1.6) ×10^−6^ s^−1^, respectively. Targets 2 and 5 were inhibited only five- and two-fold with cleavage rates of 1.87 (±0.55) ×10^−3^ s^−1^ and 5.9 (±1.7) ×10^−3^ s^−1^, respectively. The smaller inhibition at 37 °C than at 25 °C can be explained by increased nucleosome unwrapping at higher temperatures^50, 51^. We conclude that the nucleosome wrap broadly inhibits RNA-guided nucleases and that this inhibition is strongly dependent on the proximity of the target DNA to the nucleosome dyad^21–24^.

### Nucleosome Breathing Increases Cas12a Cleavage Efficiency

Our kinetic results are consistent with the hypothesis that transient DNA-histone unwrapping limits Cas12a cleavage rates. The effects of nucleosome breathing would be most pronounced at the edges of the nucleosome wrap (targets 2 & 5) and would be nearly absent for the highest-affinity histone-DNA interactions (nucleosome dyad; targets 3 & 4). We tested this hypothesis by measuring Cas12a cleavage of 601 variants that vary in nucleosome wrap tightness^46^ (**Figure 1E**, **Figure S1E**). Tight wrapping of the 601 DNA around the histone octamer is due in part to a series of ‘TA’ dinucleotides spaced 10 bp apart (asterisks in **Figure 1A**)^45, 46^. We reasoned that adding ‘TA’ dinucleotides would decrease Cas12a cleavage rates of local DNA and removing ‘TA’ dinucleotides would increase cleavage rates.

Removing the ‘TA’ repeats from the tightly wrapped left side of the 601 sequence (601ΔTA) increased the cleavage rate of target 2 five-fold, approaching the cleavage rate of opposite edge target 5 (F**igure 1E**). Conversely, adding additional ‘TA’ repeats to the inner right quartile (601RTA) reduced the target 5 cleavage rate by six-fold and also increased the target 2 cleavage rate by approximately three-fold, so that both edge targets were cleaved at the same rate. Flipping the inner half sequence centered around the dyad (601MF) increased the target 2 cleavage rate by nearly two-fold and decreased the target 5 cleavage rate by four-fold. The observed changes in Cas12a cleavage rates of the edge targets reflect the changes in flexibility of the adjacent, interior sequences. These changes in cleavage efficiency were qualitatively consistent with previous findings that 601RTA DNA is equally likely to unwrap from either side of the nucleosome when under tension and the edge-specific unwrapping probabilities of the 601MF DNA are opposite from those of the original 601 substrate^46^. We note that all 601 variants change the sequence of the nucleosome inner wrap, yet large cleavage rate changes are evident at the edge of the outer wraps. These results show that the unwrapping dynamics of the nucleosome DNA, and therefore cleavage by Cas12a, are influenced by mechanical properties of the histone octamer-DNA interactions. Additionally, our measured cleavage rates demonstrate that increased unwrapping in one half of the positioning sequence leads to tightening of the opposite DNA—a phenomenon documented previously^46, 52, 53^.

Histone post translational modifications (PTMs) also alter nucleosome unwrapping dynamics^54^. Phosphorylation of H3Y41 and acetylation of H3K56, associated with increased DNA metabolism, both increase nucleosomal DNA accessibility by increasing the unwrapping dynamics^36, 54, 55^ (highlighted in **Figure 1F**). We performed Cas12a cleavage reactions on a reconstituted human nucleosome containing amino acid substitutions in H3 (Y41E, K56Q), which are experimentally-verified mimics of these PTMs^55^. Strikingly, the nucleosome edge targets 2 and 5 were cleaved 21-fold and 3-fold faster than in the wild type nucleosome, respectively, within five-fold of their associated DNA cleavage rates (**Figure 1F**). The large, asymmetric increase in the cleavage rates of histone PTM mimetics suggests that histone PTMs that alter nucleosome stability will impact DNA accessibility to Cas nucleases^36^. Targets 3 and 4 remain refractory to cleavage within the H3(Y41E, K56Q) nucleosome, indicating that the inner dyad is inaccessible at 25 °C even within a more breathable DNA wrap. As expected, the cleavage rates for targets 1 and 6, away from the nucleosome, were identical to those for the wt nucleosome. Collectively, these data show that nucleosomal DNA cleavage is sensitive to both DNA sequence alterations and histone modifications. We conclude that nucleosomal DNA accessibility (when transiently unwrapped) is a key regulator of Cas12a cleavage efficiency.

### Nucleosomes Inhibit PAM recognition and R-loop propagation

Next, we determined whether DNA binding by Cas12a is inhibited by the nucleosome. We previously showed that R-loop formation—the second DNA binding step—is rate limiting for cleavage for short (60-nt) substrates. Thus, the cleavage rates with subsaturating concentrations of Cas12a report on the two-step DNA binding process, and the maximal cleavage rate with saturating Cas12a concentration is a direct measure of the R-loop propagation rate^8^.

We measured the DNA binding rates by performing cleavage reactions with Cas12a concentrations from 5 – 400 nM (**Figure 2A**). From the concentration-dependent cleavage rates, we determined the second-order rate constant (*k*_max_/*K*_1/2_;*k*_max_, rate of R-loop formation; *K*_1/2_, PAM binding affinity), which represents the overall rate constant for the two-step DNA binding process. Even without the nucleosome present, the binding rate constants for the four targets vary by two orders of magnitude (**Table S4**). With the nucleosome, Cas12a binding to the edge targets 2 and 5 was substantially inhibited; the rate constant decreased for target 2 binding by 320-fold to 205 (± 15) M^−1^ s^−1^, and for target 5 binding by at least 1,100-fold to 5.45 (±0.45) ×10^3^ M^−1^ s^−1^. As expected, the flanking targets 1 and 6 were relatively insensitive to the nucleosome, with the *k*_max_/*K*_1/2_ values changing less than two-fold (to 2.200 (±0.029) and 0.245 (±0.010) ×10^5^ M^−1^ s^−1^, respectively).

**Figure 2.**
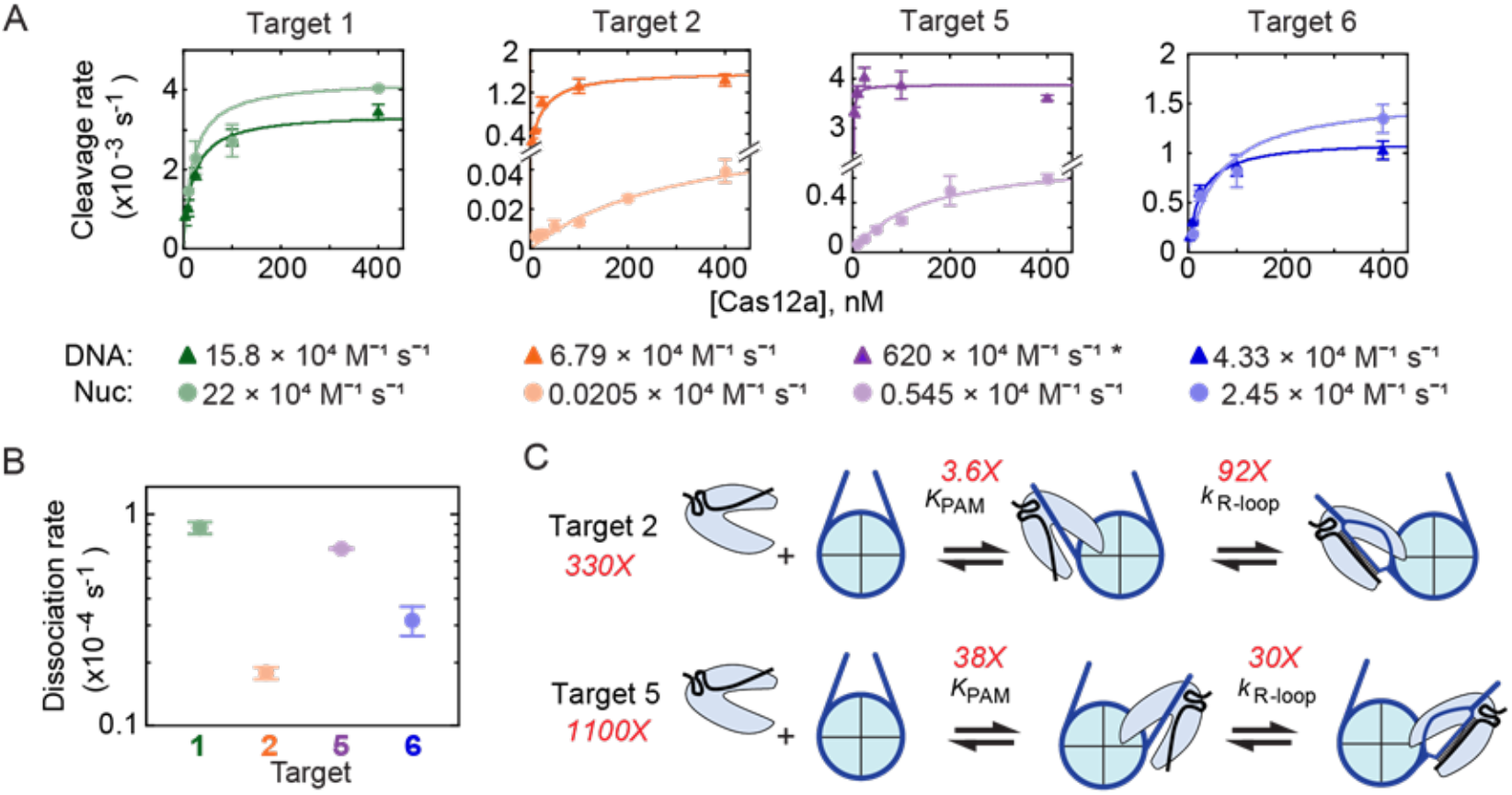
Nucleosomes inhibit Casl2a’s two-step DNA binding. A) Concentration-dependent Cas12a cleavage plots of DNA (darker colors) and nucleosome substrates (lighter colors). Colored lines: data points were fit to a hyperbolic curve. Second-order rate constants for each target and substrate are written below the associated plots. The asterisk indicates that the second-order rate constant for DNA cleavage of target 5 is a lower limit due to lack of data points in the concentration-dependent regime. B) Dissociation rate estimates. A and B) Each data point is the mean of three replicates, and error bars represent the SEM. C) Diagrams showing the measured nucleosomal inhibition on Cas12a PAM recognition and R-loop formation. *K*_PAM_: Cas12a affinity for the targets’ PAM sequence decreases due to the nucleosome; *k*_R-loop_: rate of R-loop formation decreases due to the nucleosome.

To test whether Cas12a binding remains rate limiting for DNA cleavage in the presence of a nucleosome, we measured dissociation of nuclease-dead Cas12a at these targets. We found that dissociation required hours and was much slower than DNA cleavage from the bound complex (**Figure S1A**, **2B**, **S2**). Thus, DNA binding remains rate-limiting for cleavage, and the second-order rate constants for cleavage reflect the binding kinetics both in the presence and absence of the nucleosome. Therefore, the presence of a bound nucleosome greatly decreases the Cas12a binding rate, consistent with a model of restricted access to Cas12a. Interestingly, the Cas12a dissociation rates were not necessarily faster for targets positioned within the nucleosome (**Figure 2B**).

Next, we dissected the decreased Cas12a binding rate for targets 2 and 5. The decreases in second-order rate constant noted above could result from weakened binding to the PAM sequence, slower R-loop formation, or both. To determine the effect on R-loop formation, we focused on the maximal DNA cleavage rate. This maximal rate reports directly on the rate of R-loop formation (6.6 (±2.1) and 77.4 (±6.7) ×10^−5^ s^−1^ for target 2 and 5, respectively), as cleavage of both DNA strands is expected to be much faster^8^. In absence of the nucleosome the maximal rate constants are at least 10-fold larger, indicating inhibition of R-loop formation by the nucleosome. Further, this rate constant modestly underestimates the inhibition because the cleavage reaction on DNA substrates becomes limited by the slower cleavage of the target strand at high Cas12a concentrations^8^. Comparison with the R-loop formation rate as directly measured from non-target strand (NTS) cleavage on oligonucleotide substrates (**Figure S2C**) reveals that the nucleosome slows R-loop formation for targets 2 and 5 by 92- and 30-fold, respectively. The larger inhibition for R-loop propagation at target 2 is consistent with the nucleosome entry region being more tightly wrapped than the exit region. A comparison of the *K*1/2 values in the presence of or absence of nucleosomes shows that PAM recognition is also inhibited by the bound nucleosome, by 3.6-fold and 38-fold for targets 2 and 5, respectively (**Figure 2C**). Combining the effects measured for each binding step individually gives values of 330 (±200)-fold and 1,100 (±200)-fold for inhibition to target 2 and 5, respectively (**Figure 2C**), the same within error as determined from the second-order rate constants. While the overall inhibition of binding to the two sites is similar, we speculate that a larger fraction of the inhibitory effect is focused on PAM recognition for target 5 because it is positioned closer to the nucleosome core particle.

Together our results show that the nucleosome inhibits both PAM recognition and R-loop formation, even beyond the edge of the nucleosome core particle (**Figure 2C**). This inhibition likely arises from the steric clashes between the similarly sized nucleosome and Cas12a RNP particles. Indeed, PAM recognition and R-loop propagation occur within a narrow protein channel^56, 57^ and the orientation of Cas12a’s edge targets within our substrate force the REC and NUC Lobes to point towards and overlap with the nucleosome core-creating a steric clash. Thus, nucleosome unwrapping dynamics will dominate both PAM recognition and R-loop propagation rates for targets that are close to the nucleosome.

### Proximally Targeted Cas Nucleases Trap Unwrapped Nucleosomes

Cas nuclease cleavage efficiency in cells can be enhanced by co-targeting dCas9 in close proximity to the intended genomic cut site (termed proxy-CRISPR)^58^. The biochemical basis for this enhanced cleavage is unknown. We reasoned that dCas9 captures nucleosomal DNA while it is transiently unwrapped, thereby increasing the accessibility of the proximal DNA substrate for the nuclease-active Cas12a. To directly test this model, we targeted *Streptococcus pyogenes* dCas9 (**Figure S1A**) to sequences with the canonical 3’-GGN PAM adjacent to the four Cas12a targets that either fully or partially overlap with the nucleosome (**Figure 3A**, **Figure S3**). dCas9 RNPs targeting a single proxy site (p2, p5, or p5*) were pre-incubated with the nucleosome substrate for 30 minutes before adding Cas12a RNP to begin the cleavage reaction (**Figure 3B**). Targets 3 and 4 remained uncleaved over 24 hrs when dCas9 was directed to any of the adjacent or flanking sites (**Figure S3B**), showing that dCas9 is not able to disassemble or shift the nucleosome away from its dyad.

**Figure 3.**
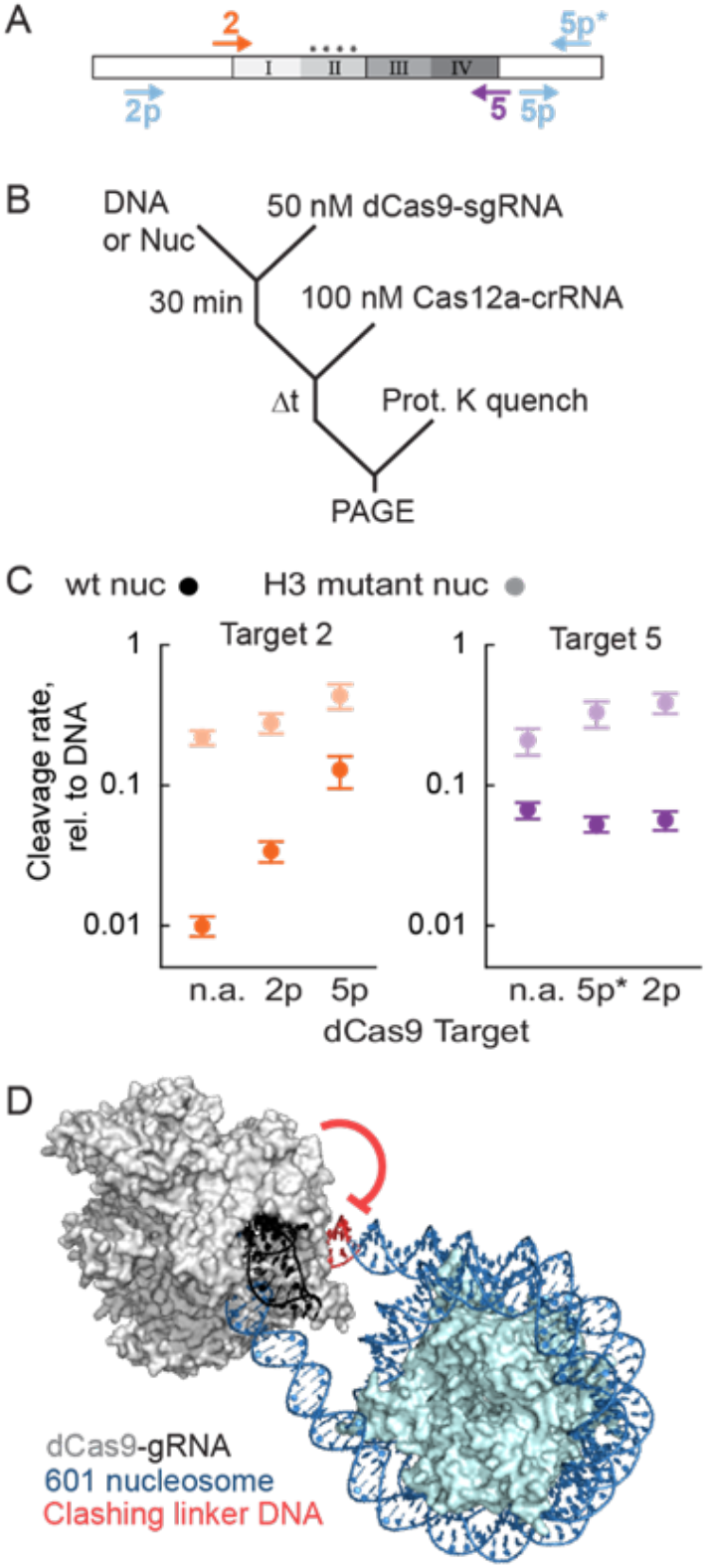
Steric interference by a proximal nuclease enhances nucleosomal DNA cleavage. A) Diagram depicting the target locations for dCas9 binding (light blue arrows) and Cas12a cleavage (color-coded arrows, same as Figure 1A). Only targets used in the main text are included here. B) Reaction scheme depicting proxy-CRISPR experimental setup. C) Proxy-CRISPR cleavage rates at the indicated dCas9 and Cas12a targets. Data points represent nucleosome cleavage by Cas12a with dCas9 pre-bound at targets 2p, 5p, 5p*, or without (n.a.) pre-bound dCas9. Data is normalized to the DNA cleavage rate at each site. Each data point is the mean of three replicates. Error bars represent the SEM. D) *SpCas9* (PDB: 4UN3^82^) modeled to bind target 5p, 13 bp away from the nucleosome (PDB: 3LZ0^81^). Linker DNA was generated by using sequence to structure modeling. Extension of the distal linker DNA shows an expected steric clash with the bound Cas9.

Target 2 cleavage rates increased more than three-fold when dCas9 was pre-bound to an adjacent sequence (target 2p, 24 bp away, **Figure 3C** orange circles), consistent with the model that nucleosome breathing dynamics dictate cleavage rates. In contrast, dCas9 targeted to 5p inhibited Cas12a cleavage of target 5 on the DNA substrate-presumably because of a steric clash between the two RNPs (six bp separate the two PAMs; **Figure S3A**). When dCas9 targets 5p* (18 bp away from target 5), the Cas12a cleavage rate for nucleosomal target 5 did not change (**Figure 3C** purple circles, **Table S6**). Target 5 is within the more unwrapped nucleosome edge, indicating that a proximal dCas9 does not increase accessibility further and may in fact have a deleterious effect on cleavage when the two targets are too close. We speculate that because target 2 is more tightly wrapped than target 5, pre-binding dCas9 peels the DNA from the histone octamer, favoring an unwrapped state. Increasing intrinsic nucleosome breathing via H3(Y41E, K56Q) mutations reduces the cleavage enhancements observed with dCas9 at target 2 (**Figure 3C**). These results highlight that intrinsic nucleosome unwrapping dynamics can substitute for a forced steric clash between dCas9 and the histone octamer.

Unexpectedly, we also see a 12-fold enhancement in target 2 cleavage by Cas12a when dCas9 is pre-bound to the distal target 5p. The stronger effect of the distal proxy-CRISPR pair suggests that cleavage enhancement may also arise due to steric clashes between the DNA-bound dCas9 and the DNA that emerges from the alternate end of the nucleosome (**Figure 3D**). We propose that binding dCas9 near the nucleosome core particle, whose large sizes are comparable, causes local DNA to peel away from the nucleosome core, but also creates steric interference for the flanking DNA at the opposite edge^59^—preventing the DNA from occupying its normal range of conformational space observed during (un)wrapping and favor more unwrapped states. Our proxy-CRISPR results further highlight the importance of nucleosome unwrapping as a key regulator of DNA cleavage by Cas12a and likely other Cas nucleases. Additionally, these experiments provide another approach to altering nucleosome unwrapping dynamics.

### DNA Accessibility is Minimally Impacted by Chromatin Phase Separation

Chromatin in the cell is organized as arrays of regularly spaced nucleosomes separated by 10 to 50-bp of linker DNA^30, 31^. Interactions between the histone tails of neighboring nucleosomes in such an array can induce liquid-liquid phase separation of nucleosomal arrays, forming a chromatin compartment^41^. We determined how Cas12a cleavage kinetics of the linker DNA are altered in nucleosome arrays under phase-separated conditions. The DNA substrate consists of 12 repeats of the 601 positioning sequence separated by 46-53 bp unique linker sequences between each nucleosome pair (**Figure 4A**). We measured Cas12a cleavage rates for two linker targets and one nucleosome-edge target (**Figure 4B**, **Table S7**). The linkers between the fourth and fifth nucleosomes (L4/5) and the eighth and ninth nucleosomes (L8/9) have a Cas12a target that is 9-13 bp away from the 601 sequence; there is sufficient space for Cas12a to bind entirely within the linker^5^. The fifth nucleosome edge target, Ex5, has a 12 bp overlap with the more unwrapped edge of the 601 sequence. We measured the cleavage rates for these targets on both DNA and reconstituted nucleosome arrays under highly dense cell-like phase-separated conditions. Cleavage reaction time points from at least three replicates were resolved on 1% agarose gels stained with Ethidium Bromide (**Figure 4D**).

**Figure 4.**
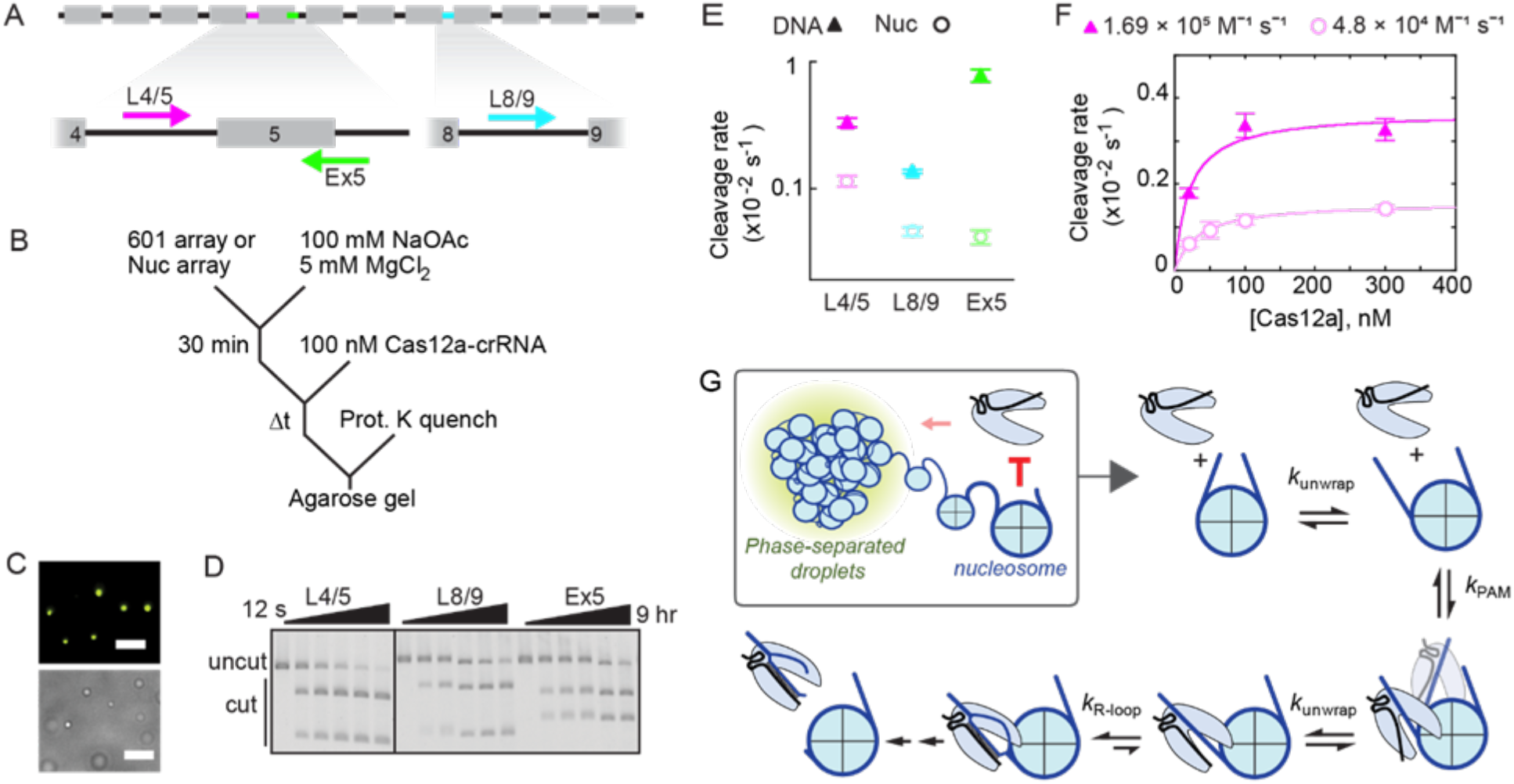
Phase-separated nucleosome arrays minimally inhibit DNA accessibility. A) Diagram of a 12-mer 601 array. Grey boxes are 601 positioning sequences. Black lines connecting the 601 repeats are unique linker DNAs. Cas12a targets are highlighted (L4/5, magenta; Ex5, green; L8/9, blue) and the associated arrows represent the direction of R-loop formation. B) Reaction scheme depicting Cas12a cleavage of DNA arrays or phase-separated nucleosome arrays. C) Fluorescent (top) and brightfield (bottom) images of phase-separated nucleosome array droplets prior to Cas12a cleavage. H2B is labeled with Alexa488 dye. Scale bars: 10 μm. D) Ethidium bromide-stained agarose gel showing cleavage products of phase-separated nucleosome arrays targeted by Cas12a. All cleavage reactions are from the same gel, and the black vertical line represents cropping of the gel. E) Cas12a cleavage plot of 601 (DNA-only) and nucleosome arrays. Cleavage of each target was slower within phase-separated nucleosome arrays. Data points are color coordinated with the targets shown in panel (A). F) Cas12a concentration-dependent cleavage plot of L4/5, targeting 601 DNA and nucleosome arrays, color coordinated as in panel (E). E and F) Each data point represents the mean of at least three replicates. Error bars represent the SEM. DNA cleavage reactions at 30 and 200 nM Cas12a were done in duplicate. G) Model of Cas12a cleavage inhibition by the nucleosome and chromatin. Chromatin condensation has a small impact on Cas12a cleavage efficiency, but the nucleosome is the key inhibitor of Cas12a by restricting its binding.

Cleavage rates for all three targets on nucleosome-free DNA substrates range from 1-8 ×10^−3^ s^−1^ with physiological concentrations of mono- and divalent salts (**Figure 4E**). We next measured the cleavage rates within reconstituted nucleosome arrays under identical buffer conditions. The nucleosome arrays form liquid-like compartments under these conditions, as shown previously^41^ and confirmed via bright field and fluorescence microscopy (**Figure 4C**). Cleavage of linker targets L4/5 and L8/9 at 100 nM Cas12a is inhibited three-fold (**Figure 4E**). Target Ex5 shows 18-fold cleavage inhibition—similar to the inhibition of target 5 positioned closely within the mono nucleosome, however this effect combines the inhibition of R-loop propagation into the nucleosome and the impact of chromatin-associated condensation. To better characterize the inhibition on target search by phase separation, we focused on linker target L4/5, measuring cleavage rates with variable Cas12a concentrations on both 601 DNA and nucleosome array substrates (**Figure 4F**). We note that the cleavage time courses of phase-separated nucleosome arrays at lower Cas12a concentrations show biphasic behavior (**Figure S4**), but due to the small amplitude of the second phase, all reactions are fit by a single exponential. The resulting second-order rate constants are 1.69 (±0.32) ×10^5^ M^−1^ s^−1^ and 4.8 (±1.4) ×10^4^ M^−1^ s^−1^, demonstrating a four-fold inhibition on Cas12a target search upon condensation of chromatin-like substrates. Our previous fluorescence recovery after photobleaching results^41^ and the penetration of TFs and chromatin modifiers within chromatin droplets suggest that Cas12a’s reduced cleavage rates are unlikely to arise from exclusion from the droplet. We also observe a reduced cleavage rate of L4/5 at saturating Cas12a concentrations, from 3.67 (±0.25) to 1.57 (±0.11) ×10^−3^ s^−1^. These results show that the nucleosome-free linker DNA remains remarkably accessible to Cas nucleases even in liquid condensates where chromatin is organized at high cell-like densities.

## Discussion

Here, we use quantitative kinetics to show that nucleosome breathing regulates cleavage by Cas12a. The nucleosome inhibits DNA cleavage by blocking both PAM recognition and R-loop propagation (**Figure 4G**). At 25 °C, Cas12a binding is inhibited by three orders of magnitude at nucleosome edges and the inner nucleosome wrap is completely inaccessible. At 37°C, cleavage within the inner wrap is inhibited three orders of magnitude relative to nucleosome-free DNA and edge targets were inhibited less than five-fold. Multiple studies have characterized how mismatches within the R-loop can reduce Cas9 and Cas12a binding and cleavage several orders of magnitude *in vitro*^8, 60–69^. Here we show that the nucleosome is a more potent barrier to DNA binding than nearly all of these partially matched targets. Our results thus explain the observation that off-target cleavage in nucleosome-depleted regions is more likely to occur than on-target cleavage within a nucleosome in cells^16, 18^. More broadly, these results highlight the importance of nucleosome dynamics in base editing, CRISPRi, and other strategies that exploit targeted DNA binding by dCas9 and dCas12a.

Nucleosome unwrapping dynamics are the key determinant of Cas12a cleavage kinetics. Increasing nucleosome unwrapping via changes to the DNA flexibility, histone modifications, or proximal dCas9 binding all increase corresponding Cas12a cleavage rates. Our results are broadly consistent with previous studies that show strong inhibition of Cas9 cleavage within the nucleosome^21–24^. Additionally, we demonstrate that PAM recognition can be hindered beyond the canonical nucleosome core. Greater unwrapping towards the nucleosome dyad is necessary for targets positioned more centrally within the nucleosome, and conversely, as targets are positioned farther away from the nucleosome, the importance of unwrapping is reduced (shown by the faded Cas12a in **Figure 4G**). Unwrapping can be promoted by a proximally directed dCas9, even when it is not in direct contact with the nucleosome: dCas9 can increase nucleosome breathing by altering the mechanical properties of the adjacent edge DNA as well as by sterically clashing with the distal linker of a nucleosome substrate. Our results provide a mechanistic underpinning for increased efficiency in *in vivo* Cas9 double-targeting strategies, such as proxy-CRISPR^58, 70^.

In addition to class 2 nucleases, the Type I CRISPR-Cascade RNA-guided effector complexes are increasingly repurposed for mammalian genome editing^71–73^. These CRISPR systems also recognize targets via a two-step binding mechanism. We speculate that the nucleosome will inhibit both Type I effectors complexes by blocking PAM recognition and R-loop propagation^21–24^. Notably, Type I-family CRISPR systems include Cas3, a processive ATP-dependent nuclease/helicase, to destroy foreign DNA. Future studies will be required to probe how Cascade binds nucleosomal DNA and how the Cascade/Cas3 complex navigates through this roadblock^74, 75^.

Finally, we observed that Cas12a can rapidly cleave condensed, chromatin-like DNA that is compartmentalized in a phase-separated droplet. Surprisingly, chromatin droplets only inhibit target cleavage four-fold relative to non-nucleosomal DNA. The modest inhibition observed for Cas12a following condensation of nucleosomal arrays (albeit without the linker histone H1^41^ or HP1^76^) could be explained by altered diffusion within the phase separated ‘droplets’ and slower two-step binding. Our results are consistent with a prior study showing that restriction enzymes are variably inhibited by Mg^2+^-dependent compaction of 17-mer nucleosome arrays^77^. Rapid diffusion of DNA-binding proteins in crowded environments have also been computationally predicted^78, 79^ and observed experimentally^32, 80^. Future studies will need to elucidate how dense, heterogenous, and target-poor chromatin may further effect CRISPR-Cas cleavage kinetics following condensation. In sum, our work highlights that dynamics of the nucleosome core particle, rather than higher-order chromatin-like organization, is the key regulator of Cas12a cleavage efficiency in cells.

## Materials and Methods

### Nucleic Acids and Proteins

All DNA oligos and GeneBlocks were purchased from IDT (**Table S1**). CRISPR RNA (crRNA) for Cas12a and guide RNA (sgRNA) for Cas9 were purchased from Synthego (**Table S2**). The mono-nucleosome 601 substrate was amplified with primers that include a 5’-Cy5 for fluorescence imaging. Unincorporated primers were removed using the Zymo DNA Clean and Concentrator-5 kit. Oligonucleotides used to measure NTS cleavage were 5’-radiolabeled with [γ–^32^P]ATP (Perkin-Elmer) as described previously^8^.

*As*Cas12a and nuclease dead *As*Cas12a (dCas12a) and *SpCas9* (dCas9) were expressed and purified as described previously^8, 63^.

Nucleosomes for mono-nucleosome experiments were reconstituted via salt dialysis^48^. Briefly, 15 μl reactions containing 200 ng/ μl Cy5-labeled 601 DNAs were mixed with human histone octamer at varying octamer:DNA ratios, typically 1.3:1 to 2:1, in 2 M NaCl. Reconstitution reactions were transferred to dialysis buttons (10,000 MWCO, Millipore) equilibrated in High Salt Reconstitution Buffer (10 mM Tris-HCl, pH 7.5, 2 M NaCl, 1 mM EDTA, pH 8.0, 1 mM DTT). Reconstitutions were done at 4 °C using a one-way pump in which No Salt Reconstitution Buffer (10 mM Tris-HCl, pH 7.5, 1 mM EDTA, pH 8.0, 1 mM DTT) was flowed into the dialysis buffer until the dialysis buttons were at 250 mM NaCl. Dialyzed reactions were then further dialyzed against 500 ml TCS Buffer (20 mM Tris-HCl, pH 7.5, 1 mM EDTA, pH 8.0, 1 mM DTT, 0.5% NP-40) for one hour and then deaggregated by centrifugation at 4 °C, max speed. The fraction of Cy5-labeled 601 DNA incorporated into nucleosomes was determined by native PAGE. Nucleosome arrays with fluorescent H2B were reconstituted as described previously^41^.

### Ribonucleoprotein complex (RNP) Assembly

Cas12a RNPs were assembled prior to each cleavage experiment using purified Cas12a and a precursor crRNA (pre-crRNA) that contained the full direct repeat sequence and a guide sequence complementary to one of the six targets within the 601 DNA substrate. Assembly reactions were performed by incubating Cas12a with a two-fold excess of crRNA for 30 min at 25 °C in 50 mM Tris-HCl, pH 8.0, 100 mM NaCl, 5 mM MgCl2, and 2 mM DTT. dCas9 RNPs were assembled using the same protocol. For nucleosome array experiments, NaCl was switched to NaOAc to promote visible phase separation^41^.

### Cas12a Cleavage Kinetics

Cleavage reactions were performed in the same buffer conditions as the assembly reaction. All cleavage reactions were initiated by adding the Cy5-labeled 601 DNA (or nucleosome) to the assembled Cas12a RNP at 25 °C. Reactions included 1-2 nM DNA and 100 nM Cas12a RNP unless otherwise indicated. At various time points, 10 μl of the 70 μl cleavage reaction were withdrawn and stopped in 5 μl of quench solution (30% glycerol, 100 mM EDTA, pH 8.0, 0.02% SDS, 2.4 μg/ μl Proteinase K).

Quenched samples were incubated at 52 °C for one hr to ensure protein digestion by Proteinase K (Thermo), resolved on a 10% native gel (0.5X TBE), and scanned using a Typhoon FLA 9500 (GE Healthcare). Bands of fluorescence corresponding to substrate and cleaved product were quantified using ImageQuant 5.2 (GE Healthcare). DNA cleavage rates were the same regardless of which end of the 601 DNA substrate was labeled with Cy5, indicating that the fluorophore did not affect DNA cleavage by Cas12a.

Cleavage reactions of the short oligo duplexes (**Table S3**) were performed as described previously^8^ but with buffer conditions altered to better match the buffer conditions of the mono-nucleosome substrate reactions: 50 mM Tris, pH 8.0, 100 mM NaCl, 5 mM MgCl_2_, 2 mM DTT, 0.2 mg/ml BSA. Quenched samples were resolved over a 20% denaturing polyacrylamide gel containing 7 M urea. Gels were exposed, imaged, and analyzed as previously described^8^.

### Mono nucleosome Dissociation Experiments

dCas12a-crRNA complexes were incubated with reconstituted nucleosomes for two hrs, (21 hrs for target 2) to promote maximum binding at 25 °C in the same buffer conditions as the cleavage reactions. The dCas12a protein includes a RuvC nuclease-inactivating point mutation (D908A), preventing DNA cleavage^56^. Pre-binding was done at above the K_1/2_ values calculated from concentration-dependent cleavage reactions: 360 nM Cas12a for targets 1, 5, and 6 and 500 nM for target 2. Reactions were diluted to 100 nM Cas12a, and Cas12a dissociation was made irreversible by addition of an oligonucleotide target strand complementary to the crRNA (**Table S3**). Two time points were sampled, mixed with pre-chilled loading buffer, put on ice-cold water, and immediately loaded onto a pre-chilled native 5% polyacrylamide gel in 0.5X TBE supplemented with 5 mM MgCl_2_ and 5% glycerol to stabilize complexes and prevent disassembly in the gel (especially important for edge targets). The native gel was maintained at 4 °C during the run to prevent complex dissociation in the gel. The signal that migrated above ‘nucleosome-only’ and ‘DNA-Cas12a’ bands was considered part of the nucleosome-bound complex and was normalized to a ‘no binding’ control lane in which nucleosome substrate and chase were pre-mixed before addition of dCas12a (representing t=∞).

### Cleavage of Phase-Separated Nucleosome Arrays

Nucleosome arrays were incubated in Phase Separation Buffer (50 mM Tris-HCl, pH 8.0, 100 mM NaOAc, 5 mM MgCl2, 2 mM DTT, 0.1 mg/ml BSA, 5% glycerol) at 1.4X the final nucleosome concentration for 30 min at 25 °C. This pre-incubation step was also included for DNA-only substrates. After pre-incubation, reactions were initiated by addition of 0.4X volumes of pre-assembled Cas12a to the DNA substrate, bringing the nucleosome final concentration to 100 nM. At various time points, 6 μl aliquots were withdrawn and stopped in 3 μl of quench solution (as defined above for reactions with mono-nucleosomes). Following digestion with Proteinase K at 52 °C, samples were run on a 1% agarose gel and stained with Ethidium Bromide. Gels were scanned using a Typhoon FLA 9500 (GE Healthcare) using fluorescence excitation and emission wavelengths of 532 nm and 600 nm, respectively. Bands of fluorescence corresponding to substrate and cleaved product were quantified using ImageQuant 5.2 (GE Healthcare).

### Fluorescence Microscopy of Phase-Separated Arrays

Nucleosome arrays with AlexaFluor 488-labeled histone H2B were prepared as described above, injected into a house-made passivated flow cell, and immediately visualized on an Eclipse Ti Nikon camera. Imaging buffer was the same buffer used in cleavage reactions. Bright-field and epifluorescence images were illuminated with a SOLA-SE-II light source (Lumencor). Data were collected with 200 ms exposure every second through a 60X water immersion objective (Nikon). Images were adjusted (contrast, coloration) in ImageJ 1.47v.

### Quantification and Statistical Analysis

Cleavage reactions for each target were performed at least three times at the indicated Cas12a concentration. The progress curves were fit by a single exponential function to determine the observed rate constant, and the reported values reflect the average value from these fits ± SEM (standard error of the mean). All data fitting was performed with Kaleidagraph. If reconstituted nucleosomes had a non-negligible amount of unincorporated Cy5-labeled DNA (maximum 20%), nucleosomal cleavage data was fit by a double exponential function in which the cleaved product corresponding to unincorporated DNA was constrained by previously determined rate and amplitude. To determine values of *k*_max_ and K1/2, the dependence of the observed rate constant on Cas12a concentration was analyzed by performing weighted data fit by a hyperbolic curve. The reported values of *k*_max_ and K_1/2_ reflect the derived values ± error of the fit. Cas12a dissociation progress curves were fit by a decreasing single exponential function, and the rate constants reported reflect the average ± standard error of the mean from three replicate experiments.

## Author Contributions

I.S., F.A.S., R.R., and I.J.F. designed the research. I.S., F.S.A., and B.A.G. performed experiments. I.S. and F.A.S. analyzed the data. I.S. and I.J.F. wrote the first draft and all authors contributed to editing the manuscript. The authors declare no conflict of interest.

## Acknowledgments

We thank members of the Finkelstein and Russell labs for helpful discussions and reagents. This work was supported by: (i) NIGMS grants P01GM066275 and R35GM131777 (to R.R.) and R01GM124141 (to I.J.F.); (ii) the Welch Foundation grants F-1563 (to R.R.), F-1808 (to I.J.F.), and I-1544 (to M.K.R.); and (iii) the Allen Foundation Distinguished Investigator Award (to M.K.R.).

## References

1. Pickar-Oliver, A. & Gersbach, C.A. The next generation of CRISPR-Cas technologies and applications. Nat Rev Mol Cell Biol 20, 490–507 (2019).

2. Swarts, D.C. & Jinek, M. Cas9 versus Cas12a/Cpf1: Structurefunction comparisons and implications for genome editing. Wiley Interdiscip Rev RNA, e1481 (2018).

3. Shmakov, S. et al. Diversity and evolution of class 2 CRISPR-Cas systems. Nat Rev Microbiol 15, 169–182 (2017).

4. Nishimasu, H. & Nureki, O. Structures and mechanisms of CRISPR RNA-guided effector nucleases. Curr Opin Struct Biol 43, 68–78 (2017).

5. Swarts, D.C., van der Oost, J. & Jinek, M. Structural Basis for Guide RNA Processing and Seed-Dependent DNA Targeting by CRISPR-Cas12a. Mol Cell 66, 221–233.e224 (2017).

6. Gong, S., Yu, H.H., Johnson, K.A. & Taylor, D.W. DNA Unwinding Is the Primary Determinant of CRISPR-Cas9 Activity. Cell Rep 22, 359–371 (2018).

7. Raper, A.T., Stephenson, A.A. & Suo, Z. Functional Insights Revealed by the Kinetic Mechanism of CRISPR/Cas9. J Am Chem Soc 140, 2971–2984 (2018).

8. Strohkendl, I., Saifuddin, F.A., Rybarski, J.R., Finkelstein, I.J. & Russell, R. in Mol Cell, Vol. 71 816–824 e813 (2018 Elsevier Inc, United States; 2018).

9. van Aelst, K., Martínez-Santiago, C.J., Cross, S.J. & Szczelkun, M.D. The Effect of DNA Topology on Observed Rates of R-Loop Formation and DNA Strand Cleavage by CRISPR Cas12a. Genes (Basel) 10 (2019).

10. Kim, D. et al. Genome-wide analysis reveals specificities of Cpf1 endonucleases in human cells. Nature biotechnology 34, 863–868 (2016).

11. Kleinstiver, B.P. et al. Genome-wide specificities of CRISPR-Cas Cpf1 nucleases in human cells. Nature biotechnology 34, 869–874 (2016).

12. Eslami-Mossallam, B. et al. A mechanistic model improves off-target predictions and reveals the physical basis of SpCas9 fidelity. biorxiv (2020).

13. Tsai, S.Q. et al. GUIDE-seq enables genome-wide profiling of off-target cleavage by CRISPR-Cas nucleases. Nat Biotechnol 33, 187–197 (2015).

14. Kim, D. et al. Digenome-seq: genome-wide profiling of CRISPR-Cas9 off-target effects in human cells. Nat Methods 12, 237–243, 231 p following 243 (2015).

15. Pattanayak, V. et al. High-throughput profiling of off-target DNA cleavage reveals RNA-programmed Cas9 nuclease specificity. Nat Biotechnol 31, 839–843 (2013).

16. Wu, X. et al. Genome-wide binding of the CRISPR endonuclease Cas9 in mammalian cells. Nat Biotechnol 32, 670–676 (2014).

17. Chari, R., Mali, P., Moosburner, M. & Church, G.M. Unraveling CRISPR-Cas9 genome engineering parameters via a library-on-library approach. Nat Methods 12, 823–826 (2015).

18. Kuscu, C., Arslan, S., Singh, R., Thorpe, J. & Adli, M. Genome-wide analysis reveals characteristics of off-target sites bound by the Cas9 endonuclease. Nat Biotechnol 32, 677–683 (2014).

19. Knight, S.C. et al. Dynamics of CRISPR-Cas9 genome interrogation in living cells. Science 350, 823–826 (2015).

20. Verkuijl, S.A. & Rots, M.G. The influence of eukaryotic chromatin state on CRISPR-Cas9 editing efficiencies. Curr Opin Biotechnol 55, 68–73 (2019).

21. Horlbeck, M.A. et al. Nucleosomes impede Cas9 access to DNA in vivo and in vitro. eLife 5 (2016).

22. Yarrington, R.M., Verma, S., Schwartz, S., Trautman, J.K. & Carroll, D. Nucleosomes inhibit target cleavage by CRISPR-Cas9 in vivo. Proc Natl Acad Sci U S A 115, 9351–9358 (2018).

23. Isaac, R.S. et al. Nucleosome breathing and remodeling constrain CRISPR-Cas9 function. eLife 5 (2016).

24. Hinz, J.M., Laughery, M.F. & Wyrick, J.J. Nucleosomes Inhibit Cas9 Endonuclease Activity in Vitro. Biochemistry 54, 7063–7066 (2015).

25. Luger, K., Mader, A.W., Richmond, R.K., Sargent, D.F. & Richmond, T.J. Crystal structure of the nucleosome core particle at 2.8 A resolution. Nature 389, 251–260 (1997).

26. Andrews, A.J. & Luger, K. Nucleosome structure(s) and stability: variations on a theme. Annu Rev Biophys 40, 99–117 (2011).

27. Cutter, A.R. & Hayes, J.J. A brief review of nucleosome structure. FEBS Lett 589, 2914–2922 (2015).

28. McKnight, S.L. & Miller, O.L. Ultrastructural patterns of RNA synthesis during early embryogenesis of Drosophila melanogaster. Cell 8, 305–319 (1976).

29. Woodcock, C.L., Safer, J.P. & Stanchfield, J.E. Structural repeating units in chromatin. I. Evidence for their general occurrence. Exp Cell Res 97, 101–110 (1976).

30. Szerlong, H.J. & Hansen, J.C. Nucleosome distribution and linker DNA: connecting nuclear function to dynamic chromatin structure. Biochem Cell Biol 89, 24–34 (2011).

31. Valouev, A. et al. Determinants of nucleosome organization in primary human cells. Nature 474, 516–520 (2011).

32. Fierz, B. & Poirier, M.G. Biophysics of Chromatin Dynamics. Annu Rev Biophys 48, 321–345 (2019).

33. Polach, K.J. & Widom, J. Mechanism of protein access to specific DNA sequences in chromatin: a dynamic equilibrium model for gene regulation. J Mol Biol 254, 130–149 (1995).

34. Li, G. & Widom, J. Nucleosomes facilitate their own invasion. Nat Struct Mol Biol 11, 763–769 (2004).

35. Anderson, J.D., Thastrom, A. & Widom, J. Spontaneous access of proteins to buried nucleosomal DNA target sites occurs via a mechanism that is distinct from nucleosome translocation. Mol Cell Biol 22, 7147–7157 (2002).

36. North, J.A. et al. Regulation of the nucleosome unwrapping rate controls DNA accessibility. Nucleic Acids Res 40, 10215–10227 (2012).

37. Tims, H.S., Gurunathan, K., Levitus, M. & Widom, J. Dynamics of nucleosome invasion by DNA binding proteins. J Mol Biol 411, 430–448 (2011).

38. Anderson, J.D. & Widom, J. Sequence and position-dependence of the equilibrium accessibility of nucleosomal DNA target sites. J Mol Biol 296, 979–987 (2000).

39. Luo, Y., North, J.A., Rose, S.D. & Poirier, M.G. Nucleosomes accelerate transcription factor dissociation. Nucleic Acids Res 42, 3017–3027 (2014).

40. Gebala, M., Johnson, S.L., Narlikar, G.J. & Herschlag, D. Ion counting demonstrates a high electrostatic field generated by the nucleosome. Elife 8 (2019).

41. Gibson, B.A. et al. Organization of Chromatin by Intrinsic and Regulated Phase Separation. Cell (2019).

42. Larson, A.G. et al. Liquid droplet formation by HP1α suggests a role for phase separation in heterochromatin. Nature 547, 236–240 (2017).

43. Li, G., Levitus, M., Bustamante, C. & Widom, J. Rapid spontaneous accessibility of nucleosomal DNA. Nat Struct Mol Biol 12, 46–53 (2005).

44. Winogradoff, D. & Aksimentiev, A. Molecular Mechanism of Spontaneous Nucleosome Unraveling. J Mol Biol 431, 323–335 (2019).

45. Lowary, P.T. & Widom, J. New DNA sequence rules for high affinity binding to histone octamer and sequence-directed nucleosome positioning. J Mol Biol 276, 19–42 (1998).

46. Ngo, T.T., Zhang, Q., Zhou, R., Yodh, J.G. & Ha, T. Asymmetric unwrapping of nucleosomes under tension directed by DNA local flexibility. Cell 160, 1135–1144 (2015).

47. Hinz, J.M., Laughery, M.F. & Wyrick, J.J. Nucleosomes Selectively Inhibit Cas9 Off-target Activity at a Site Located at the Nucleosome Edge. J Biol Chem 291, 24851–24856 (2016).

48. Dyer, P.N. et al. Reconstitution of nucleosome core particles from recombinant histones and DNA. Methods Enzymol 375, 23–44 (2004).

49. Bondarenko, V.A. et al. in Mol Cell, Vol. 24 469–479 (United States; 2006).

50. Blossey, R. & Schiessel, H. The dynamics of the nucleosome: thermal effects, external forces and ATP. Febs j 278, 3619–3632 (2011).

51. Chereji, R.V. & Morozov, A.V. Functional roles of nucleosome stability and dynamics. Brief Funct Genomics 14, 50–60 (2015).

52. Konrad, S.F. High-throughput AFM analysis reveals unwrapping pathways of H3 and CENP-A nucleosomes. biorxiv.org (2020).

53. Bilokapic, S., Strauss, M. & Halic, M. Histone octamer rearranges to adapt to DNA unwrapping. Nat Struct Mol Biol 25, 101–108 (2018).

54. Bowman, G.D. & Poirier, M.G. Post-translational modifications of histones that influence nucleosome dynamics. Chem Rev 115, 2274–2295 (2015).

55. Brehove, M. et al. Histone core phosphorylation regulates DNA accessibility. J Biol Chem 290, 22612–22621 (2015).

56. Yamano, T. et al. Crystal Structure of Cpf1 in Complex with Guide RNA and Target DNA. Cell 165, 949–962 (2016).

57. Gao, P., Yang, H., Rajashankar, K.R., Huang, Z. & Patel, D.J. Type V CRISPR-Cas Cpf1 endonuclease employs a unique mechanism for crRNA-mediated target DNA recognition. Cell Res 26, 901–913 (2016).

58. Chen, F. et al. Targeted activation of diverse CRISPR-Cas systems for mammalian genome editing via proximal CRISPR targeting. Nat Commun 8, 14958 (2017).

59. Bednar, J. et al. Structure and Dynamics of a 197 bp Nucleosome in Complex with Linker Histone H1. Mol Cell 66, 384–397.e388 (2017).

60. Singh, D., Sternberg, S.H., Fei, J., Doudna, J.A. & Ha, T. Real-time observation of DNA recognition and rejection by the RNA-guided endonuclease Cas9. Nat Commun 7, 12778 (2016).

61. Singh, D. et al. Real-time observation of DNA target interrogation and product release by the RNA-guided endonuclease CRISPR Cpf1 (Cas12a). Proc Natl Acad Sci U S A 115, 5444–5449 (2018).

62. Boyle, E.A. et al. High-throughput biochemical profiling reveals sequence determinants of dCas9 off-target binding and unbinding. Proc Natl Acad Sci U S A 114, 5461–5466 (2017).

63. Jr, S.K.J. et al. (biorxiv; 2019).

64. Sternberg, S.H., Redding, S., Jinek, M., Greene, E.C. & Doudna, J.A. DNA interrogation by the CRISPR RNA-guided endonuclease Cas9. Nature 507, 62–67 (2014).

65. Ivanov, I.E. et al. Cas9 interrogates DNA in discrete steps modulated by mismatches and supercoiling. Proc Natl Acad Sci U S A 117, 5853–5860 (2020).

66. Szczelkun, M.D. et al. Direct observation of R-loop formation by single RNA-guided Cas9 and Cascade effector complexes. Proc Natl Acad Sci U S A 111, 9798–9803 (2014).

67. Murugan, K., Seetharam, A.S., Severin, A.J. & Sashital, D.G. CRISPR-Cas12a has widespread off-target and dsDNA-nicking effects. J Biol Chem 295, 5538–5553 (2020).

68. Fu, B.X.H. et al. Target-dependent nickase activities of the CRISPR-Cas nucleases Cpf1 and Cas9. Nat Microbiol 4, 888–897 (2019).

69. Singh, D. et al. Mechanisms of improved specificity of engineered Cas9s revealed by single-molecule FRET analysis. Nat Struct Mol Biol 25, 347–354 (2018).

70. Bolukbasi, M.F. et al. Orthogonal Cas9-Cas9 chimeras provide a versatile platform for genome editing. Nat Commun 9, 4856 (2018).

71. Cameron, P. et al. Harnessing type I CRISPR-Cas systems for genome engineering in human cells. Nat Biotechnol 37, 1471–1477 (2019).

72. Pickar-Oliver, A. et al. Targeted transcriptional modulation with type I CRISPR-Cas systems in human cells. Nat Biotechnol 37, 1493–1501 (2019).

73. Dolan, A.E. et al. Introducing a Spectrum of Long-Range Genomic Deletions in Human Embryonic Stem Cells Using Type I CRISPR-Cas. Mol Cell 74, 936–950.e935 (2019).

74. Dillard, K.E. et al. Assembly and Translocation of a CRISPR-Cas Primed Acquisition Complex. Cell 175, 934–946.e915 (2018).

75. Loeff, L., Brouns, S.J.J. & Joo, C. Repetitive DNA Reeling by the Cascade-Cas3 Complex in Nucleotide Unwinding Steps. Mol Cell 70, 385–394.e383 (2018).

76. Sanulli, S. et al. HP1 reshapes nucleosome core to promote phase separation of heterochromatin. Nature 575, 390–394 (2019).

77. Poirier, M.G., Bussiek, M., Langowski, J. & Widom, J. Spontaneous access to DNA target sites in folded chromatin fibers. J Mol Biol 379, 772–786 (2008).

78. Krepel, D., Gomez, D., Klumpp, S. & Levy, Y. Mechanism of Facilitated Diffusion during a DNA Search in Crowded Environments. J Phys Chem B 120, 11113–11122 (2016).

79. Kanada, R., Terakawa, T., Kenzaki, H. & Takada, S. Nucleosome Crowding in Chromatin Slows the Diffusion but Can Promote Target Search of Proteins. Biophys J 116, 2285–2295 (2019).

80. Hihara, S. et al. Local nucleosome dynamics facilitate chromatin accessibility in living mammalian cells. Cell Rep 2, 1645–1656 (2012).

81. Vasudevan, D., Chua, E.Y.D. & Davey, C.A. Crystal structures of nucleosome core particles containing the ‘601’ strong positioning sequence. J Mol Biol 403, 1–10 (2010).

82. Anders, C., Niewoehner, O., Duerst, A. & Jinek, M. Structural basis of PAM-dependent target DNA recognition by the Cas9 endonuclease. Nature 513, 569–573 (2014).

