## Supplemental Figures and Tables for "Inhibition of CRISPR-Cas12a DNA Targeting by Nucleosomes and Chromatin"

Isabel Strohkendl[1,*], Fatema A. Saifuddin[1,2], Bryan A. Gibson[3], Michael K. Rosen[3], Rick Russell[1,*], Ilya J. Finkelstein[1,2,*]

[1]Department of Molecular Biosciences and Institute for Cellular and Molecular Biology, University of Texas at Austin, Austin, Texas 78712, USA

[2]Center for Systems and Synthetic Biology, University of Texas at Austin, Austin, Texas 78712, USA

[3]Department of Biophysics and Howard Hughes Medical Institute, University of Texas Southwestern Medical Center, Dallas, TX 75390, USA

[1]To whom correspondence should be addressed:
Isabel Strohkendl:
Rick Russell:
Ilya J. Finkelstein:

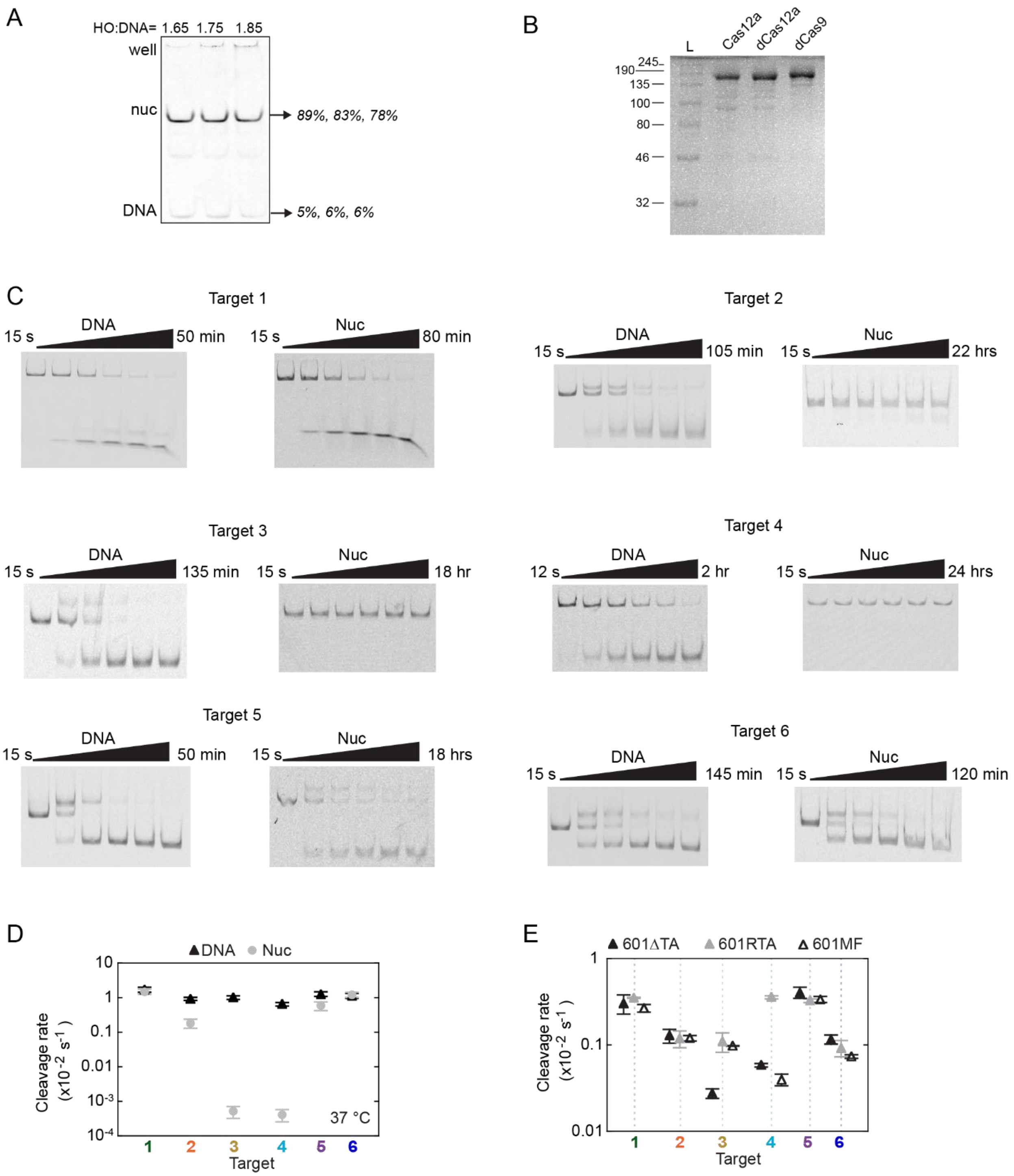


**Figure S1. Mono-nucleosome inhibits Cas12a cleavage.**

A) Native PAGE gel showing an example of a nucleosome reconstitution titration. Histone octamers (HO) are added in increasing amounts relative to DNA molarity. Percentages to the right report on the main nucleosome band signal intensity. B) SDS PAGE of purified CRISPR-Cas proteins used in kinetic assays. C) Representative Cas12a cleavage gels of DNA and nucleosome targets at 25°C. We attribute the appearance of a slower migrating species to be nicked DNA resulting from NTS cleavage. D) Cleavage plot of Cas12a targeting DNA or nucleosomes at 37 °C. Except for the higher temperature, reactions were performed exactly as those shown in Figure 1C. E) Cleavage plot of Cas12a targeting DNA-only substrates of the 601 sequence variants at 25 °C. D and E) Data points represent the average of at least three replicates. Error bars represent the SEM.

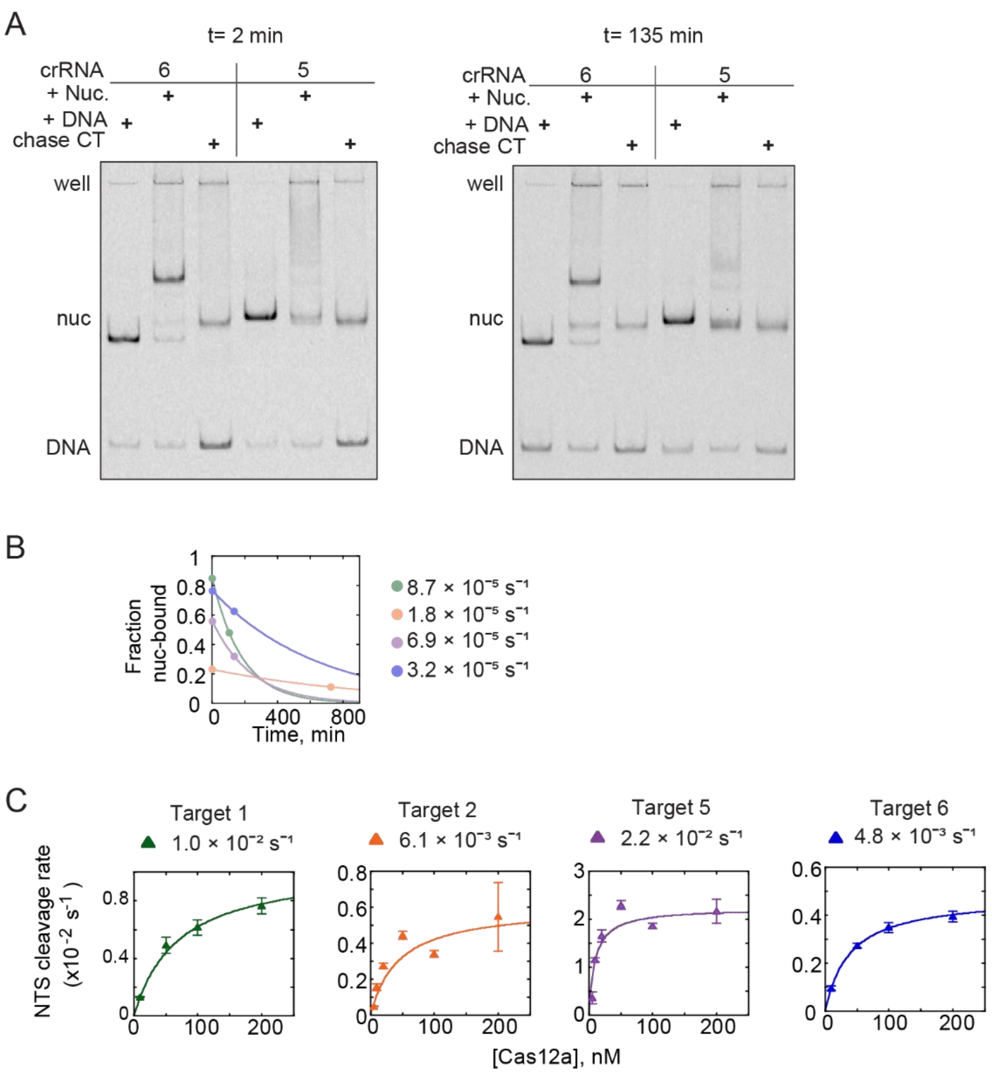


**Figure S2. Nucleosomes inhibit Cas12a's two-step binding.**

A) Example native gels used to determine dissociation rate estimates of Cas12a from nucleosome substrates at 2 and 135 minutes after beginning the dissociation experiment. Here, nuclease dead dCas12a is pre-bound to either the nucleosome or DNA by crRNA complementary to targets 5 or 6. The 'chase CT' lanes confirm unlabeled, complementary target strand chase blocks binding of Cas12a to the nucleosome and DNA and is used as the time course end point. Signal representing a super shift (band for target 6, smear for target 5) was quantified as nucleosome-bound (not including the well). B) Example dissociation curves using two sampled time points and an endpoint determined by the chase control. The low starting value for target 2 is due to the relatively close rates for binding and dissociation within the given reaction conditions. Rates are color-coded as in Figure 2A. C) Concentration-dependent Cas12a cleavage plots of the nontarget strand (NTS) of a ~60 bp oligo duplex. The $k_{max}$ value (shown) was used as a reporter for unimpeded R-loop formation. Each data point is the mean of at least three replicates and error bars represent the SEM.

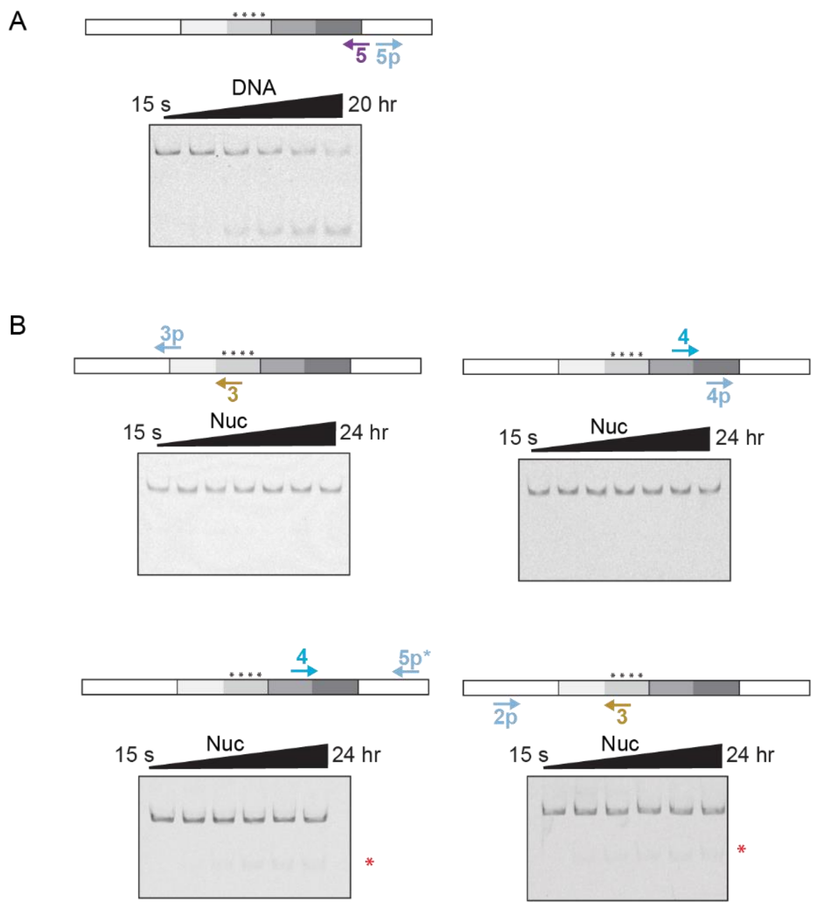


**Figure S3. proxy-CRISPR cleavage of inner wrap Cas12a targets.**

A) proxy-cleavage gel showing 100-fold inhibition of target 5 cleavage by Cas12a when Cas9 is pre-bound adjacent at target 5p. B) Diagrams depicting proxy-CRISPR pairs for Cas12a targets within the inner wrap of the nucleosome that did not lead to enhanced nucleosome cleavage. Below are the example gels of the related cleavage reaction. The red asterisks mark small bursts of cleavage products that are accounted for by a small fraction Cy5-DNA that was unincorporated into a nucleosome after the reconstitution, with the observed rate matching target-specific DNA cleavage rates. The two cleavage reactions on the left were performed with the same batch of reconstituted nucleosomes, and the two to the right from a second batch.

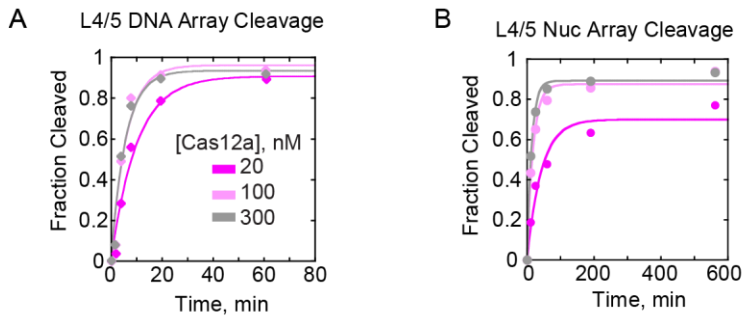


**Figure S4. Concentration-dependent cleavage of linker L4/5.**

Time courses depicting L4/5 cleavage of A) 601 DNA-only array and B) a phase-separated nucleosome array. All cleavage rates were determined by fitting to a single exponential, however we note a double exponential curve is the better fitting for nucleosome array cleavage.

| DNA | DNA Sequence (5′ to 3′) |
|---|---|
| 601 | CTGCTAGATCACAGACTCCAGCCAGAACTGTTCATCCTTAAAATCCCTTATGTGATGGACCCTATTTA<br>TGACTACC**CTGGAGAATCCCGGTGCCGAGGCCGCTCAATTGGTCGTAGACAGCTCTAGCACCGCTTAA**<br>**ACGCACGTACGCGCTGTCCCCCGCGTTTTAACCGCCAAGGGGATTACTCCCTAGTCTCCAGGCACGTG**<br>**TCAGATATATACATCCTGT**GCGTAAATTGAATCCAGCGTCTCATCTTTATGCGTCTAAAGAGATCGGA<br>AGAGCG |
| 601MF | CTGCTAGATCACAGACTCCAGCCAGAACTGTTCATCCTTAAAATCCCTTATGTGATGGACCCTATTTA<br>TGACTACC**CTGGAGAATCCCGGTGCCGAGGCCGCTCAATTGCTAGGGAGTAATCCCCTTGGCGGTTAA**<br>**AACGCGGGGGACACCGCGTACGTGCGTTTAAGCGGTGCTAGAGCTGTCTACGACTCTCCAGGCACGTG**<br>**TCAGATATATACATCCTGT**GCGTAAATTGAATCCAGCGTCTCATCTTTATGCGTCTAAAGAGATCGGA<br>AGAGCG |
| 601ΔTA | CTGCTAGATCACAGACTCCAGCCAGAACTGTTCATCCTTAAAATCCCTTATGTGATGGACCCTATTTA<br>TGACTACC**CTGGAGAATCCCGGTGCCGAGGCCGCTCAATTGGTCGccGACAGCTCggGCACCGCTTAA**<br>**ACGCACGccCGCGCTGTCCCCCGCGTTTTAACCGCCAAGGGGATTACTCCCTAGTCTCCAGGCACGTG**<br>**TCAGATATATACATCCTGT**GCGTAAATTGAATCCAGCGTCTCATCTTTATGCGTCTAAAGAGATCGGA<br>AGAGCG |
| 601RTA | CTGCTAGATCACAGACTCCAGCCAGAACTGTTCATCCTTAAAATCCCTTATGTGATGGACCCTATTTA<br>TGACTACC**CTGGAGAATCCCGGTGCCGAGGCCGCTCAATTGGTCGTAGACAGCTCTAGCACCGCTTAA**<br>**ACGCACGTACGCGCTGTCTACCGCGTTTTAACCGCCAATAGGATTACTTACTAGTCTCCAGGCACGTG**<br>**TCAGATATATACATCCTGT**GCGTAAATTGAATCCAGCGTCTCATCTTTATGCGTCTAAAGAGATCGGA<br>AGAGCG |
| FS132 | Cy5-CGCTCTTCCGATCTCTTTAGACGCATAAAGATGAGACGCTGGA |
| FS133 | Cy5-CTGCTAGATCACAGACTCCAGCCAGAACTGTTCATCCTTAAA |
| FS134 | CGCTCTTCCGATCTCTTTAGACGCATAAAGATGAGACGCTGGA |
| FS135 | CTGCTAGATCACAGACTCCAGCCAGAACTGTTCATCCTTAAA |

**Table S1. Mono-nucleosome DNA sequences and primers.** Bold represents the variant Widom 601 nucleosome positioning sequences. We use the 601MF and 601RTA nomenclature first presented in Ngo, T *et al* 2015. However, our 601MF has several more bp flipped to avoid disrupting target 3 and 4 target sequences recognized by Cas12a.

| Target | Guide RNA Sequence (5′ to 3′) |
|---|---|
| Cas12a 1 | AGGAUGAACAGUUCUGGCUGGAGU |
| Cas12a 2 | UGACUACCCUGGAGAAUCCCGGUG |
| Cas12a 3 | AGCGGUGCUAGAGCUGUCUACGAC |
| Cas12a 3 ΔTA | AGCGGUGCCCGAGCUGUCGGCGAC |
| Cas12a 4 | ACCGCCAAGGGGAUUACUCCCUAG |
| Cas12a 4 RTA | ACCGCCAAUAGGAUUACUUACUAG |
| Cas12a 5 | CGCACAGGAUGUAUAUAUCUGACA |
| Cas12a 6 | GACGCAUAAAGAUGAGACGCUGGA |
| Cas12a L4/5 | CCACCAUGGGAAUUCCUUAUUAUA |
| Cas12a L8/9 | GAUAUGGUACCGAAUUCCGGUGUU |
| Cas12a Ex5 | ACAUCACCACAGGAUGUAUAUAUC |
| Cas9 2p | UAAGGAUGAACAGUUCUGGC |
| Cas9 3p | AUGACUACCCUGGAGAAUCC |
| Cas9 4p | GUAUAUAUCUGACACGUGCC |
| Cas9 5p | GACGCAUAAAGAUGAGACGC |
| Cas9 5p* | CUUUAUGCGUCUAAAGAGAU |

**Table S2. Table of Cas12a and Cas9 RNA guide sequences.** RNA sequences listed are complementary to targets within the mono-nucleosome substrate and the 12-mer 601 array substrate. Cas12a crRNAs and Cas9 gRNAs were ordered from Synthego with their associated direct repeat sequences. Cas9 5p sequence was positioned too close to Cas12a target 5 (Cas12a cleavage on DNA pre-bound with dCas9 was inhibited) so Cas9 target 5p* was used for adjacent proximal targeting with Cas12a 5.

| Oligo | Sequence (5′ to 3′) |
|---|---|
| Target 1 NTS | CGCTCTTCCGATCTTTTAAGGATGAACAGTTCTGGCTGGAGTGTAGCTACTGTGCT |
| Target 1 TS | AGCACAGTAGCTACACTCCAGCCAGAACTGTTCATCCTTAAAAGATCGGAAGAGCG |
| Target 2 NTS | CGCTCTTCCGATCTTTTATGACTACCCTGGAGAATCCCGGTGGTAGCTACTGTGCT |
| Target 2 TS | AGCACAGTAGCTACCACCGGGATTCTCCAGGGTAGTCATAAAAGATCGGAAGAGCG |
| Target 5 NTS | CGCTCTTCCGATCTTTTACGCACAGGATGTATATATCTGACAGTAGCTACTGTGCT |
| Target 5 TS | AGCACAGTAGCTACTGTCAGATATATACATCCTGTGCGTAAAAGATCGGAAGAGCG |
| Target 6 NTS | CGCTCTTCCGATCTTTTAGACGCATAAAGATGAGACGCTGGAGTAGCTACTGTGCT |
| Target 6 TS | AGCACAGTAGCTACTCCAGCGTCTCATCTTTATGCGTCTAAAAGATCGGAAGAGCG |

**Table S3. DNA oligonucleotide sequences for NTS cleavage rates.** Short oligo duplex DNA substrates were constructed from two fully-complementary oligonucleotides representing the nontarget strand (NTS) and the target strand (TS). Duplexes were used for determining rate of unimpeded R-loop formation in Figure 2 and Figure S2.

| Octamer | DNA | Temp. (°C) | [Cas12a] (nM) | Cas12a Target Cleavage Rates ($\times 10^{-3}$ $s^{-1}$) | | | | | |
|---|---|---|---|---|---|---|---|---|---|
| | | | | **1** | **2** | **3** | **4** | **5** | **6** |
| - | 601 | 25 | 100 | 2.78 ± 0.20 | 1.31 ± 0.14 | 0.85 ± 0.19 | 0.523 ± 0.086 | 3.88 ± 0.28 | 0.82 ± 0.16 |
| - | 601 | 37 | 100 | 17.3 ± 2.8 | 9.50 ± 0.98 | 10.3 ± 1.2 | 6.61 ± 0.83 | 12.9 ± 1.9 | 11.6 ± 1.9 |
| WT | 601 | 25 | 100 | 2.72 ± 0.40 | 0.0135 ± 0.0016 | 0.0006 | 0.0006 | 0.258 ± 0.029 | 0.82 ± 0.16 |
| WT | 601 | 37 | 100 | 15.2 ± 2.5 | 1.87 ± 0.55 | 0.0052 ± 0.0019 | 0.0042 ± 0.0016 | 5.9 ± 1.7 | 12.4 ± 2.1 |
| - | 601MF | 25 | 100 | 2.69 ± 0.28 | 0.75 ± 0.14 | 0.66 ± 0.17 | 0.468 ± 0.036 | 2.57 ± 0.39 | 0.42 ± 0.12 |
| - | 601ΔTA | 25 | 100 | 3.06 ± 0.76 | 1.29 ± 0.23 | 0.278 ± 0.036 | 0.587 ± 0.029 | 4.06 ± 0.60 | 1.17 ± 0.14 |
| - | 601RTA | 25 | 100 | 3.531 ± 0.065 | 1.19 ± 0.26 | 1.11 ± 0.27 | 3.60 ± 0.15 | 3.28 ± 0.14 | 0.93 ± 0.20 |
| WT | 601MF | 25 | 100 | 2.67 ± 0.42 | 0.0235 ± 0.0043 | 0.0006 | 0.0006 | 0.0784 ± 0.0046 | 0.389 ± 0.017 |
| WT | 601ΔTA | 25 | 100 | 2.66 ± 0.61 | 0.0661 ± 0.0030 | 0.0006 | 0.0006 | 0.163 ± 0.014 | 0.98 ± 0.10 |
| WT | 601RTA | 25 | 100 | 4.067 ± 0.083 | 0.0433 ± 0.0021 | 0.0006 | 0.0006 | 0.043 ± 0.022 | 1.36 ± 0.48 |
| Mut, H3 Y41E/K56Q | 601 | 25 | 100 | 3.06 ± 0.13 | 0.283 ± 0.017 | 0.0006 | 0.0006 | 0.83 ± 0.27 | 0.644 ± 0.053 |
| - | 601 | 25 | 5 | 0.82 ± 0.24 | 0.267 ± 0.048 | | | 3.333 ± 0.096 | 0.164 ± 0.026 |
| - | 601 | 25 | 10 | 1.02 ± 0.21 | 0.467 ± 0.019 | | | 3.833 ± 0.096 | 0.311 ± 0.015 |
| - | 601 | 25 | 25 | 1.89 ± 0.15 | 1.022 ± 0.084 | | | 4.04 ± 0.19 | 0.622 ± 0.053 |
| - | 601 | 25 | 400 | 3.44 ± 0.20 | 1.44 ± 0.12 | | | 3.611 ± 0.056 | 1.033 ± 0.092 |
| WT | 601 | 25 | 10 | 1.456 ± 0.043 | 0.0067 ± 0.0021 | | | 0.068 ± 0.015 | 0.19 ± 0.03 |
| WT | 601 | 25 | 25 | 2.29 ± 0.42 | 0.0075 ± 0.0022 | | | 0.116 ± 0.017 | 0.578 ± 0.095 |
| WT | 601 | 25 | 50 | | 0.0116 ± 0.0027 | | | 0.190 ± 0.033 | |
| WT | 601 | 25 | 200 | | 0.0256 ± 0.0015 | | | 0.5 ± 0.2 | |
| WT | 601 | 25 | 400 | 4.056 ± 0.056 | 0.0392 ± 0.0057 | | | 0.596 ± 0.066 | 1.35 ± 0.25 |

| Target | Naked Cleavage at 25°C ($\times 10^{4}$ $M^{-1}$ $s^{-1}$) | Nucleosome Cleavage at 25°C ($\times 10^{3}$ $M^{-1}$ $s^{-1}$) | Nucleosome $K_{1/2}$ (nM) | Nucleosome $k_{max}$ ($\times 10^{-3}$ $s^{-1}$) |
|---|---|---|---|---|
| 1 | 15.80 ± 0.56 | 220.0 ± 2.9 | 19.2 ± 1.0 | 4.240 ± 0.063 |
| 2 | 6.79 ± 0.16 | 0.205 ± 0.015 | 320 ± 160 | 0.066 ± 0.021 |
| 5 | 620 *(lower limit)* | 5.45 ± 0.45 | 142 ± 20 | 0.774 ± 0.067 |
| 6 | 4.33 ± 0.12 | 24.5 ± 1.0 | 64 ± 11 | 1.58 ± 0.19 |

**Table S4. Cas12a cleavage rates of human mono-nucleosome targets 1-6**. Rates are an average of at least three replicates ± SEM. Rate constants were determined by performing weighted-data fits to report fit parameter value ± error of the fit.

| **Oligoduplex Target** | **[Cas12a]** (nM) | **Cas12a NTS Cleavage Rates** ($\times 10^{-3}$ $s^{-1}$) | **Oligoduplex Target** | **[Cas12a]** (nM) | **Cas12a NTS Cleavage Rates** ($\times 10^{-3}$ $s^{-1}$) |
|---|---|---|---|---|---|
| 1 | 10 | 1.322 ± 0.029 | 6 | 10 | 0.956 ± 0.098 |
| 1 | 50 | 4.93 ± 0.56 | 6 | 50 | 2.73 ± 0.10 |
| 1 | 100 | 6.17 ± 0.53 | 6 | 100 | 3.50 ± 0.20 |
| 1 | 200 | 7.67 ± 0.56 | 6 | 200 | 3.95 ± 0.23 |
| 2 | 5 | 0.505 ± 0.029 | 5 | 5 | 3.6 ± 1.2 |
| 2 | 10 | 1.56 ± 0.18 | 5 | 10 | 11.53 ± 0.41 |
| 2 | 20 | 2.75 ± 0.14 | 5 | 20 | 16.5 ± 1.3 |
| 2 | 50 | 4.44 ± 0.23 | 5 | 50 | 23.00 ± 0.89 |
| 2 | 100 | 3.42 ± 0.20 | 5 | 100 | 18.67 ± 0.59 |
| 2 | 200 | 5.4 ± 1.9 | 5 | 200 | 21.6 ± 2.5 |
| **1** | $k_{max}$ ($\times 10^{-3}$ $s^{-1}$): 10.48 ± 0.89 | | **6** | $k_{max}$ ($\times 10^{-3}$ $s^{-1}$): 4.81 ± 0.32 | |
| **2** | $k_{max}$ ($\times 10^{-3}$ $s^{-1}$): 6.07 ± 0.42 | | **5** | $k_{max}$ ($\times 10^{-3}$ $s^{-1}$): 22.30 ± 0.71 | |

**Table S5. Cas12a concentration dependent non-target strand (NTS) cleavage rates.** Compiled rates of NTS cleavage of a short oligoduplex for targets 2 and 5. Rates are an average of at least three replicates ± SEM. Rate constants were determined by performing weighted-data fits to report fit parameter value ± error of the fit.

| Octamer | dSpCas9 Target | Cas12a Target Cleavage Rates ($\times 10^{-3}$ $s^{-1}$) 2 | 5 |
|---|---|---|---|
| WT | 2p | 0.0449 ± 0.0060 | 0.222 ± 0.031 |
| WT | 5p | 0.169 ± 0.040 | *0.206 ± 0.020 |
| Mut, H3 Y41E/K56Q | 2p | 0.367 ± 0.042 | 1.51 ± 0.22 |
| Mut, H3 Y41E/K56Q | 5p | 0.578 ± 0.098 | *1.28 ± 0.26 |

**Table S6. Table of Cas12a ‘proxy CRISPR’ cleavage rates.** Rates of Cas12a cleavage for edge Targets 2 and 5. Rates are reported per second, average of at least three replicates ± SEM. For Cas9 targets adjacent to Cas12a target 5, the Cas9 target is 5p*, rates are marked by an ‘*’.

| **Substrate** | **Phase Separation** | **[Cas12a]** (nM) | **Cas12a Target Cleavage Rates** ($\times10^{-3}$ $s^{-1}$) **L4/5** | **L8/9** | **Ex5** |
|---|---|---|---|---|---|
| DNA Array | No | 100 | 3.36 ± 0.28 | 1.38 ± 0.05 | 7.84 ± 0.932 |
| DNA Array | No | 20 | 1.79 ±.11 | | |
| DNA Array | No | 300 | 3.27 ± 0.25 | | |
| Nucleosome Array | Yes | 100 | 1.16 ± 0.11 | 0.458 ± 0.038 | 0.428 ± 0.039 |
| Nucleosome Array | Yes | 20 | 0.613 ± 0.085 | | |
| Nucleosome Array | Yes | 50 | 0.93 ± 0.19 | | |
| Nucleosome Array | Yes | 300 | 1.43 ± 0.082 | | |

| **Target L4/5** | $k_{max}$ ($\times10^{-3}$ $s^{-1}$) | **Second-order Rate Constant** ($\times10^{4}$ $M^{-1}$ $s^{-1}$) |
|---|---|---|
| DNA Array | 3.66 ± 0.25 | 16.9 ± 3.2 |
| Nucleosome Array | 1.57 ± 0.11 | 4.8 ± 1.4 |

**Table S7. Cas12a chromatin cleavage rates.** Rates of Cas12a cleavage for linker and edge targets within a 12-mer 601 array. Rates are reported per second, average of at least two replicates for DNA array and three replicates for nucleosome array ± SEM. Rate constants were determined by performing weighted-data fits to report fit parameter value ± error of the fit.
